# *in silico* Whole Genome Sequencer & Analyzer (iWGS): a computational pipeline to guide the design and analysis of *de novo* genome sequencing studies

**DOI:** 10.1101/028134

**Authors:** Xiaofan Zhou, David Peris, Jacek Kominek, Cletus P Kurtzman, Chris Todd Hittinger, Antonis Rokas

**Affiliations:** Department of Biological Sciences, Vanderbilt University, Nashville, TN 37235, USA; Laboratory of Genetics, Genome Center of Wisconsin, DOE Great Lakes Bioenergy Research Center, Wisconsin Energy Institute, J. F. Crow Institute for the Study of Evolution, University of Wisconsin-Madison, Madison, WI 53706, USA; Mycotoxin Prevention and Applied Microbiology Research Unit, National Center for Agricultural Utilization Research, Agricultural Research Service, US Department of Agriculture, Peoria, IL, USA

**Keywords:** genome sequencing, high-throughput sequencing, *de novo* assembly, experimental design, simulation, non-model organism

## Abstract

The availability of genomes across the tree of life is highly biased toward vertebrates, pathogens, human disease models, and organisms with relatively small and simple genomes. Recent progress in genomics has enabled the *de novo* decoding of the genome of virtually any organism, greatly expanding its potential for understanding the biology and evolution of the full spectrum of biodiversity. The increasing diversity of sequencing technologies, assays, and *de novo* assembly algorithms have augmented the complexity of *de novo* genome sequencing projects in non-model organisms. To reduce the costs and challenges in *de novo* genome sequencing projects and streamline their experimental design and analysis, we developed iWGS (*<u>i</u>n silico* <u>W</u>hole <u>G</u>enome <u>S</u>equencer and Analyzer), an automated pipeline for guiding the choice of appropriate sequencing strategy and assembly protocols. iWGS seamlessly integrates the four key steps of a *de novo* genome sequencing project: data generation (through simulation), data quality control, *de novo* assembly, and assembly evaluation and validation. The last three steps can also be applied to the analysis of real data. iWGS is designed to enable the user to have great flexibility in testing the range of experimental designs available for genome sequencing projects, and supports all major sequencing technologies and popular assembly tools. Three case studies illustrate how iWGS can guide the design of *de novo* genome sequencing projects and evaluate the performance of a wide variety of user-specified sequencing strategies and assembly protocols on genomes of differing architectures. iWGS, along with a detailed documentation, is freely available at https://github.com/zhouxiaofan1983/iWGS.

## Introduction

Whole genome sequences are rich sources of information about organisms that are superbly useful for addressing a wide variety of evolutionary questions, such as measuring mutation rates (Kumar and Subramanian 2002), characterizing the genomic basis of adaptation (Roux, et al. 2014), and building the tree of life (Rokas, et al. 2003; Salichos and Rokas 2013). Until now, however, organismal diversity has been highly unevenly covered, and most sequenced genomes correspond to model organisms, organisms of medical or economic importance, or ones that have relatively small and simple genomes (Reddy, et al. 2015).

The rapid advance of DNA sequencing technologies has dramatically reduced the labor and cost required for genome sequencing, which is evidenced by the burst of large-scale genome projects in recent years that includes, for example, the 1,000 Fungal Genomes (1KFG) Project (Grigoriev, et al. 2011), the Yeast 1,000 Plus (Y1000+) Project (Hittinger, et al. 2015), the Insect 5K Project (Robinson, et al. 2011), and the Genome 10K Project (Genome 10K Community of Scientists 2009). Some of these projects have already begun to fuel important discoveries in evolution and other fields (Zhang, et al. 2014). Equally importantly, high-throughput DNA sequencing has made it possible for single investigators to perform *de novo* genome sequencing in virtually any organism they are interested in (Rokas and Abbot 2009). Such sequencing efforts may target various organisms with a large diversity of genome architectures. Therefore, to achieve optimal results, the choice of sequencing strategy (i.e. the combination of sequencing technology (e.g. Illumina, Pacific Biosciences), sequencing assay (e.g. paired-end, mate-pair), and other variables, such as sequencing depth) and assembly protocols (e.g. assemblers and the associated parameters) should ideally be tailored to the characteristics of a given genome, such as size and GC/repeat content (Nagarajan and Pop 2013).

The vast majority of *de novo* sequenced genomes have been generated using the Illumina technology, either solely or in combination with other technologies (Reddy, et al. 2015). This is largely due to the Illumina technology’s ability to quickly generate tens to hundreds of millions of highly accurate short sequence reads of up to 300 bases per run at very low per base cost (Glenn 2011). Additionally, the Illumina technology offers two powerful sequencing assays, paired-end (PE) and mate-pair (MP), which generate sequence read pairs that span short (hundreds of base-pairs) and relatively long (thousands of base-pairs) genomic regions, respectively. Mixing multiple PE and MP libraries with different insert sizes allows for highly flexible sequencing strategies, and several state-of-the-art assembly algorithms have been developed that exploit all these advantages. For instance, the *de novo* genome assembler ALLPATHS-LG can generate high quality draft assemblies for mammalian-size genomes using only Illumina short-read data by including both MP and overlapping PE libraries (Gnerre, et al. 2011). On its own, however, the Illumina technology performs less well for more complex genomes, mainly due to the short lengths of Illumina sequence reads and the technology’s bias against certain genomic regions (e.g. GC-rich) (Ross, et al. 2013).

The Pacific Biosciences (PacBio) technology generates sequence reads that are substantially longer and have much less sequencing bias, albeit at the cost of a substantially lower per-read accuracy; the average read length increases to above 10 kilo-bases with the latest chemistry but displays only ∼87% accuracy (Koren and Phillippy 2015). Thus, this technology is particularly useful for the sequencing of complex genomes (Koren and Phillippy 2015). Recent developments in both sequencing chemistry and assembly algorithms have enabled PacBio-only *de novo* assembly for microbial genomes (Koren, et al. 2013), but the high sequence coverage required for this approach remains cost-prohibitive for large eukaryotic genomes. Nevertheless, in combination with more affordable Illumina short-read data, PacBio long reads – even at low coverage – can lead to significantly improved assemblies (Utturkar, et al. 2014; McIlwain, et al. 2016).

*De novo* genome sequencing projects are further complicated by the large array of assembly software tools, which differ in many aspects, such as algorithmic design, supported/required data types, and computational efficiency (Nagarajan and Pop 2013; Simpson and Pop 2015). Systematic evaluations of assembly programs show that no single assembler is the best across all circumstances; rather, an assembler’s performance critically depends on genome complexity and sequencing strategy adopted (Earl, et al. 2011; Bradnam, et al. 2013). Moreover, many assemblers use adjustable parameters (e.g. the k-mer size for *de Bruijn* assemblers), the values of which can critically affect the assembly quality. In practice, such parameters are often selected intuitively or through the time-consuming process of testing multiple values.

The great number of possible ways to combine sequencing technologies, assays, and assembly algorithms poses a great challenge for the experimental design and data analysis in *de novo* genome sequencing projects, which in turn can sometimes lead to poor quality or downright incorrect assemblies (Denton, et al. 2014). As a consequence, several pipelines have been developed to automate specific steps in the process; for example, the recently developed iMetAMOS (Koren, et al. 2014) and RAMPART (Mapleson, et al. 2015) have been specifically designed to automate genome assembly. However, as *de novo* genome sequencing is increasingly adopted by single investigator laboratories, there is an urgent need for streamlined approaches that enable investigators to not only efficiently generate high-quality draft genome assemblies but also to predict (via simulation) and identify the most suitable design(s) (i.e. the most suitable combination(s) of sequencing strategy and assembly protocol) currently available for a specific genome.

To address this need, we have developed an automated pipeline for the design and execution of *de novo* genome sequencing projects that we name iWGS (*<u>i</u>n silico* <u>W</u>hole <u>G</u>enome <u>S</u>equencer and Analyzer). To approximate the performance of different sequencing strategies and assembly protocols, iWGS simulates high-throughput genome sequencing on user-provided reference genomes (e.g. genomes that closely represent the characteristics of the real targets), facilitating the identification of optimal experimental designs. iWGS allows users to experiment with various combinations of sequencing technologies, assays, assembly tools, and relevant parameters in a single run. iWGS is also designed to work with real data and can be used as a convenient tool for automated selection of the best assembly or genome assembler. Finally, using three case studies, each one focused on specific challenges frequently encountered in *de novo* genome sequencing studies (e.g. high repeat content, biased nucleotide composition, etc.), we illustrate how iWGS can be applied to guiding the design and analysis of *de novo* genome sequencing studies.

## Results

### The design of iWGS

iWGS encompasses all major steps of a typical *de novo* genome sequencing study, including the generation of sequence reads, data quality control, *de novo* assembly, and evaluation of assemblies (Fig. 1).

**Figure 1.**
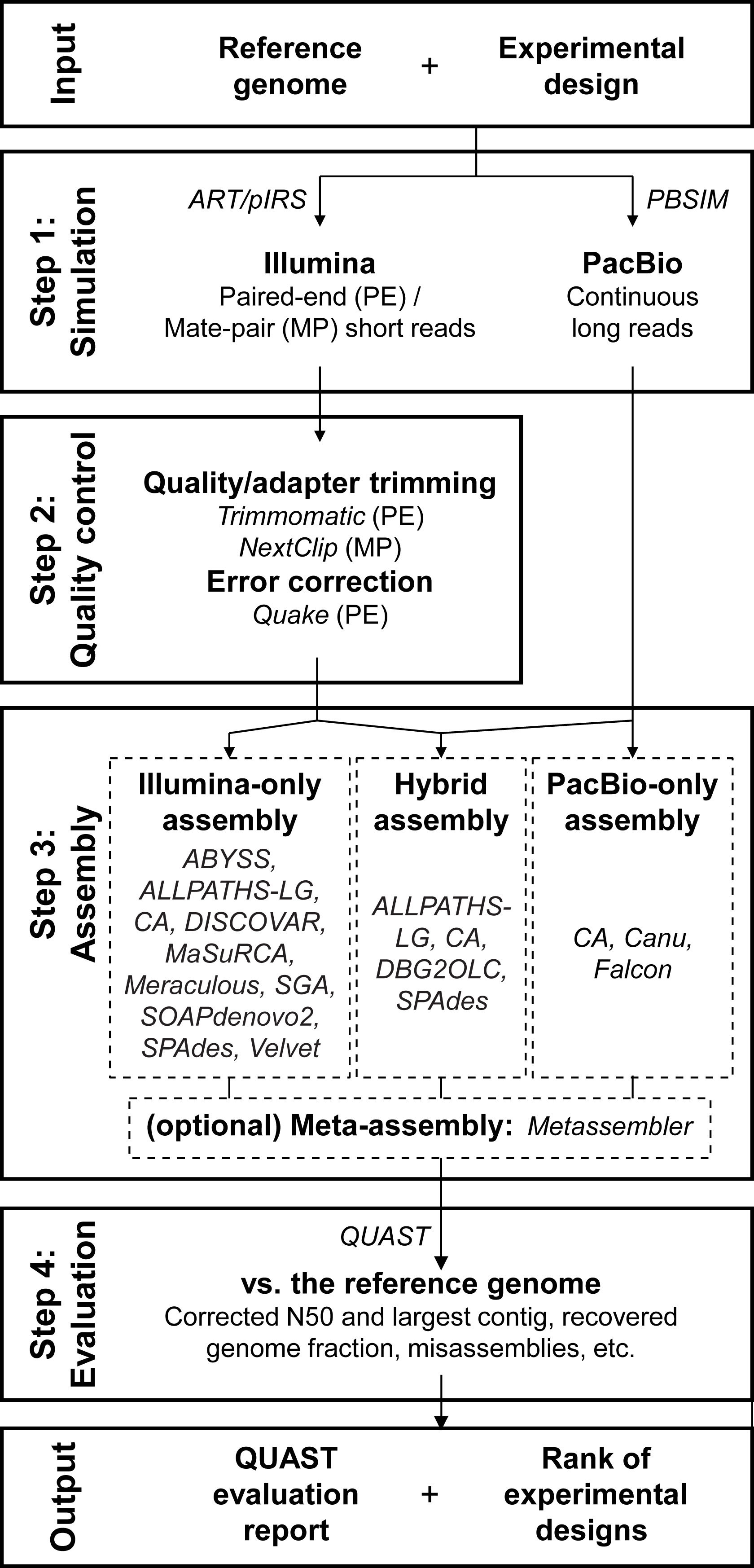
iWGS workflow. A typical iWGS analysis consists of four steps: 1) data simulation (optional); 2) preprocessing (optional); 3) *de novo* assembly; and 4) assembly evaluation. iWGS supports both Illumina short reads and PacBio long reads, and a wide selection of assemblers to enable *de novo* assembly using either or both types of data. Users can start the analysis simulating data drawn from a reference genome assembly or, alternatively, use real sequencing data as input and skip the simulation step.

1. **Simulation:** iWGS uses the realistic high-throughput sequencing (HTS) read simulators ART (Huang, et al. 2012), pIRS (Hu, et al. 2012), and PBSIM (Ono, et al. 2013) to generate Illumina and PacBio sequence reads from a given user-specified genome. These programs can simulate all popular data types, including Illumina PE and MP sequence reads, as well as PacBio continuous long sequence reads. The distributions of read quality and read length are easily adjustable for both Illumina and PacBio data. Furthermore, these simulators mimic sequencing errors and nucleotide composition biases in real data by using empirical profiles of these artifacts, which can be easily customized to stay current with upgrades in sequencing technologies. For instance, we have created a quality-score frequency profile learned from sequence reads generated by latest PacBio chemistry to better reflect the improved sequence read accuracy. This simulation step can be omitted when the goal is the analysis of real data. Alternatively, the users may choose to perform only the simulation and use the simulated data for other analyses.
2. **Quality control:** HTS data generated by all technologies contain errors and artifacts, which may sometimes substantially compromise the quality of the assembly (Zhou and Rokas 2014). Therefore, iWGS includes an optional step to perform preprocessing of the data, including trimming of low-quality bases, removal of adapter contaminations, and correction of sequencing errors. Since some assemblers [e.g. ALLPATHS-LG (Ribeiro, et al. 2012)] have their own pre-processing modules, iWGS automatically determines for each assembly protocol whether to use the original or the processed data.
3. **Assembly:** To maximize users’ flexibility in experimental design, iWGS supports 15 *de novo* genome assembly tools [ABYSS (Simpson, et al. 2009), ALLPATHS-LG (Ribeiro, et al. 2012), Celera Assembler (Myers, et al. 2000; Berlin, et al. 2015), Canu (Berlin, et al. 2015), DBG2OLC (Ye, et al. 2014), DISCOVAR (Weisenfeld, et al. 2014), Falcon (https://github.com/PacificBiosciences/falcon), MaSuRCA (Zimin, et al. 2013), Meraculous (Chapman, et al. 2011), Minia (Salikhov, et al. 2013), Platanus (Kajitani, et al. 2014), SGA(Simpson and Durbin 2012), SOAPdenovo2 (Luo, et al. 2012), SPAdes (and a diploid-aware version called dipSPAdes) (Bankevich, et al. 2012), and Velvet (Zerbino and Birney 2008)], most of which have participated in recent large-scale assembler comparisons (Bradnam, et al. 2013; Magoc, et al. 2013). These supported assemblers allow users to carry out *de novo* assembly using only Illumina short-read data (e.g. SOAPdenovo2) and only PacBio long-read data (e.g. Canu and Falcon), or to perform hybrid assembly that uses both (e.g. SPAdes and DBG2OLC). To achieve the best possible results while avoiding the computationally expensive process of testing multiple combinations of parameters, iWGS takes advantage of successful assembly recipes (i.e., recommended settings for each assembler) established in studies such as Assemblathon 2 (Bradnam, et al. 2013) and GAGE-B (Magoc, et al. 2013), and uses KmerGenie to determine the optimal k-mer size (Chikhi and Medvedev 2014). In addition, assemblies generated from different underlying data and/or assembly algorithms can be merged using Metassembler (Wences and Schatz 2015) to achieve a potentially better final assembly.
4. **Evaluation:** iWGS uses QUAST (Gurevich, et al. 2013) to evaluate all generated assemblies. In addition to providing basic statistics like N50 (the largest contig/scaffold size wherein half of the total assembly size is contained in contigs/scaffolds no shorter than this value), QUAST compares each assembly against the reference genome (in the case of simulations) and generates a number of highly informative quality matrices, such as mis-assemblies, assembled sequences not present in the reference (and vice versa), and genes recovered in the assembly if the reference genome is annotated. At the end, iWGS ranks all assemblies based on selected matrices in the QUAST report using a previously described weighting strategy (Abbas, et al. 2014). This ranking, along with the detailed QUAST report, helps users to identify the best overall assembly, as well as the corresponding combination of sequencing strategy and assembly protocol. REAPR, which utilizes the sequence data itself for assembly evaluation, is also implemented to better suit real data analysis (Hunt, et al. 2013).

iWGS is designed with flexibility and ease-of-use in mind to allow users to readily examine various experimental designs; each data set may be used multiple times in different assembly protocols, and each assembler may be run repeatedly with different input data sets. Multiple sequencing strategies and assembly protocols can be specified in a straightforward fashion in a single configuration file; only a few parameters are required for each strategy/protocol, while other settings (e.g. quality profiles for read simulation) are globally shared across strategies/protocols of the same type. Alternatively, advanced users can opt to customize the strategies/protocols so that, for example, each sequencing data set is simulated with different quality settings. Furthermore, iWGS rigorously checks the configurations for issues such as the compatibility between sequencing strategies and assembly protocols.

iWGS is a lightweight pipeline written in Perl. The source code, detailed documentation, and example test sets are freely available at https://github.com/zhouxiaofan1983/iWGS. Like many other bioinformatics pipelines, iWGS inevitably relies on a number of third-party software tools to carry out individual analyses such as data simulation and genome assembly. However, most of the tools, including at least one for each of the four major steps aforementioned, either have pre-complied executables or can be compiled locally with ease. For the convenience of users, we also include in the package scripts to automate the acquisition and installation of most software dependencies. The users can also customize the selection of tools to install according to their own needs and computational environments.

### Case Studies

To demonstrate the use of iWGS and provide examples of its utility, we developed three case studies where iWGS was used to guide the selection of sequencing strategy for genomes representing a wide range of sizes and complexity levels (Table S1). The competing strategies were selected to enable both Illumina-only and PacBio-only assemblies, as well as hybrid assembly of the two data types (Table 1). To examine the effectiveness of the simulation step of our approach, we also analyzed real sequencing data that largely match our simulation settings.

**Table 1.**
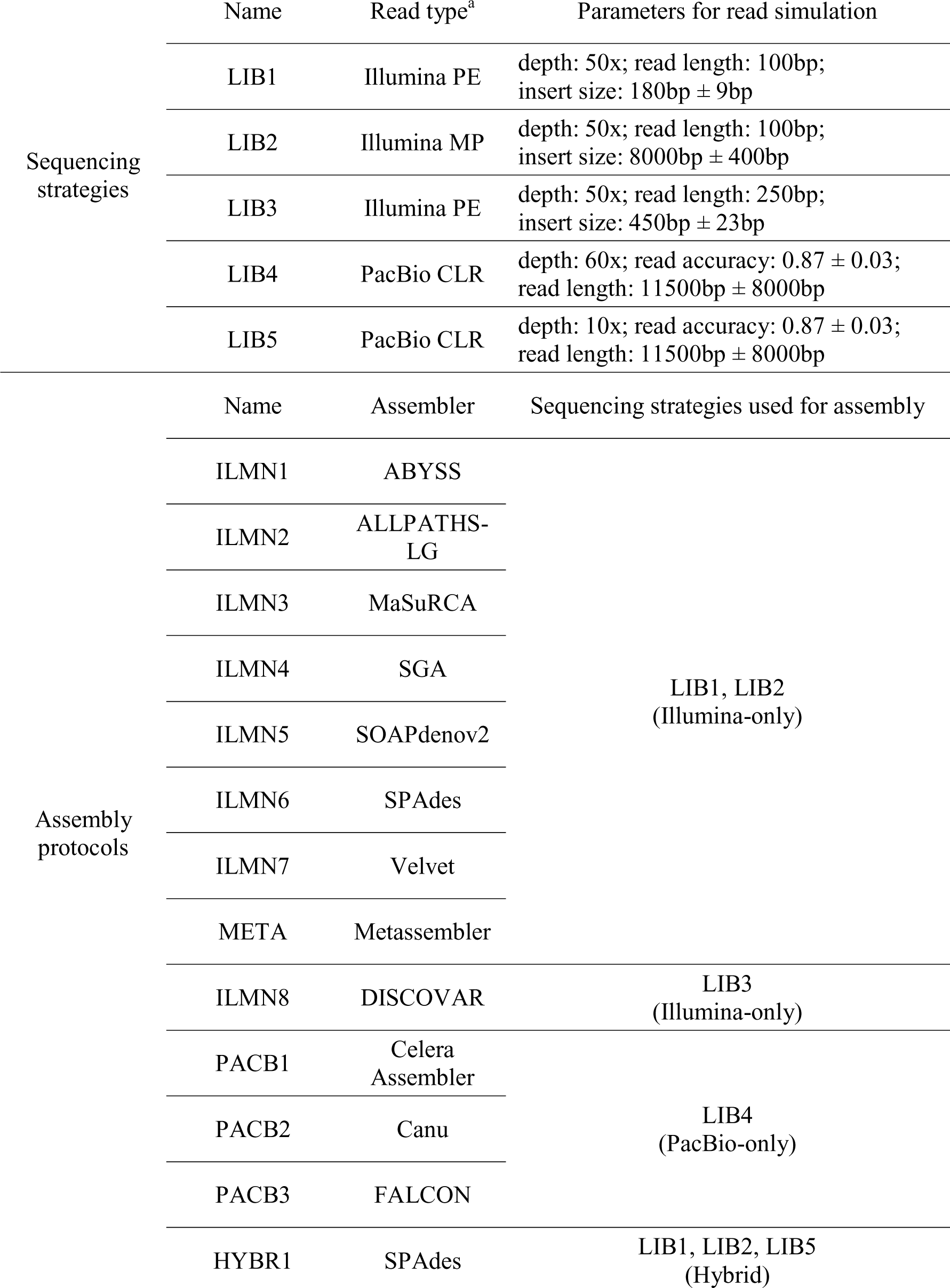
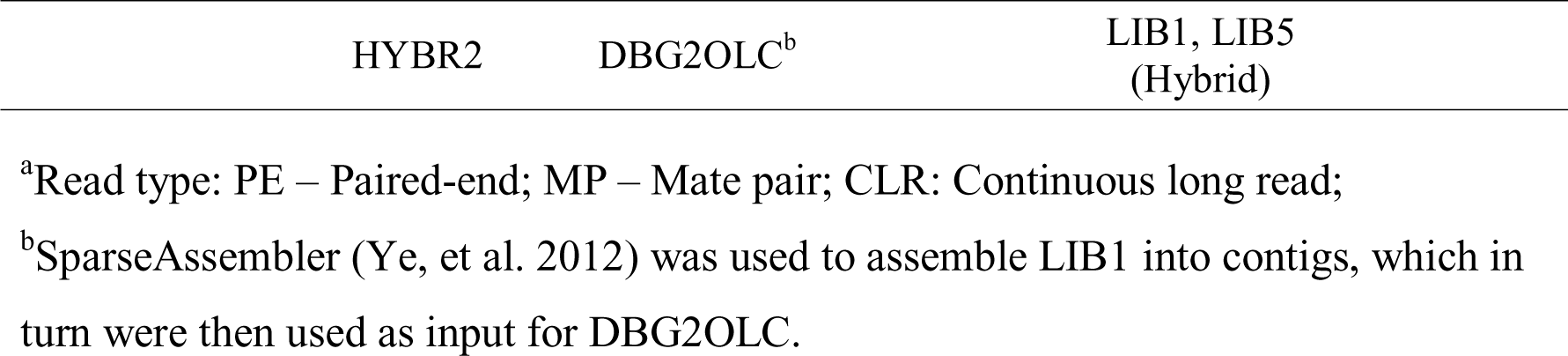
Sequencing strategies and assembly protocols evaluated in the three case studies.

*Case study I* (*Repeat-content issue*). We first compared the sequencing of two fungi, *Zymoseptoria tritici* (synonym: *Mycosphaerella graminicola*) (Goodwin, et al. 2011) and *Pseudocercospora fijiensis* (synonym: *Mycosphaerella fijiensis*) (Ohm, et al. 2012), which both belong to the class Dothideomycetes yet have dramatically different repeat contents; the estimated repeat contents are ∼15% and ∼50% for the two genomes, respectively. Our simulations showed that, while good quality assemblies can be obtained for *Z. tritici* using either data type, the PacBio-only assembly for *Ps. fijiensis* vastly outperforms assemblies based on Illumina data alone or in combination with low-coverage PacBio data (Fig. 2). The results are consistent with the notion that PacBio long reads are particularly powerful in resolving repeats (Koren, et al. 2013). We then further tested if these results are informative for guiding the sequencing of another highly repetitive Dothideomycetes genome, *Cenococcum geophilum*, which has a repeat content of c.a. 76% (http://genome.jgi.doe.gov/Cenge3). For *C. geophilum*, the PacBio-only assembly was again found to be the best, while the hybrid assembly using DBG2OLC and the Illumina-only assembly using ALLPATHS-LG were next in rank (Fig. 2; Table S2), nicely recapitulating the results of *Ps. fijiensis*. We also performed meta-assembly of Illumina-only assemblies ILMN1 to ILMN7 (Table 1) on all three genomes using Metassembler. While the meta-assembly approach substantially improved the assembly continuity for *Z. tritici*, no improvement was observed for *Ps. fijiensis* and *C. geophilum* (Fig. 2; Table S2). These results suggest that the use of iWGS would provide critical information to help end users choose a successful sequencing of highly repetitive genomes that share similar characteristics. Importantly, since simulated assemblies are recoverable, the likely impact of the different assembly strategies on genes, gene families, or pathways of interest could also be examined in detail.

**Figure 2.**
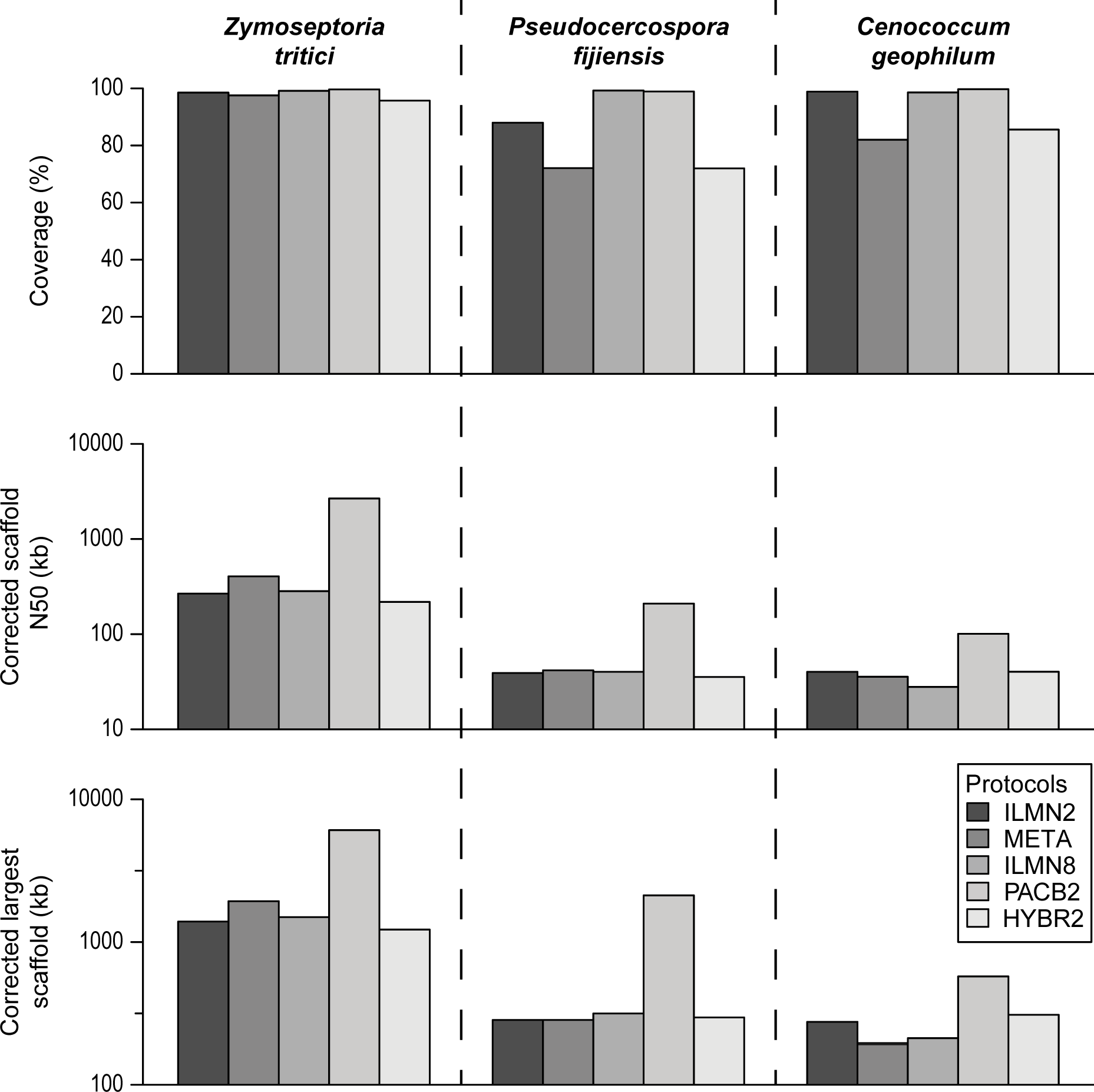
Performance comparison of five representative experimental designs on three Dothideomycetes genomes. The five designs shown include three Illumina-only designs (ILMN2: ALLPATHS-LG, META: Metassembler, and ILMN8: DISCOVAR), the best performing PacBio-only design (PACB2: Canu), and the best performing hybrid design (HYBR2: DBG2OLC) for each genome. The statistics on assembled fraction of the reference genome, scaffold N50, and largest scaffold size are all after correction for assembly errors using the reference genome as reported by QUAST in GAGE mode. By default, QUAST (in GAGE mode) corrects contigs/scaffolds by breaking them at assembly errors larger than 5bp. Scaffold N50 and largest scaffold size are shown in log10 scale.

*Case Study II* (*GC-content and mtDNA assembly issue*). We next examined the *de novo* assembly of mitochondrial genomes from whole genome sequencing data of *Saccharomyces cerevisiae* (Mewes, et al. 1997; Foury, et al. 1998). Yeast mitochondrial genomes are valuable resources for evolutionary and functional studies (Freel, et al. 2015), yet the acquisition of finished mitochondrial genome assemblies is not trivial because of their very low GC-content (∼17%). We simulated a genome sequencing experiment using the nuclear and mitochondrial genomes of *S. cerevisiae*. We tested two ratios of nuclear to mitochondrial genome copy numbers representing low (1:50) and high (1:200) mitochondrial contents respectively (Solieri 2010). iWGS analysis showed that the *S. cerevisiae* mitochondrial genome was fully recovered at both low and high mitochondrial contents using Illumina data (Table 2). Consistent with recent observations made during the assembly of the *Saccharomyces eubayanus* genome, only certain assemblers performed well; for example, ALLPATHS-LG performed surprisingly poorly, while SPAdes performed quite well (Baker, et al. 2015). Importantly, the complete mitochondrial genome can be obtained as a single contig using only Illumina or only PacBio data or using both data types (Table 2). Similarly, both Illumina and PacBio data resulted in good quality assemblies of the nuclear genome (Table S2). At the same time, different assemblers exhibited widely different performances even with the same input data (Table 2).

**Table 2.** Performance of all experimental designs evaluated in case study II.

| Nuclear:mito<br>genome ratio | Performance of strategies<sup>*</sup> |  |  |  |
| --- | --- | --- | --- | --- |
| | Complete, single<br>contig assembly<br>of the<br>mitochondrial<br>genome | Assembled<br>fraction of<br>mitochondrial<br>genome $\geq 99\%$ | 20% ≤<br>Assembled<br>fraction of<br>mitochondrial<br>genome $< 99\%$ | Assembled<br>fraction of<br>mitochondrial<br>genome $< 20\%$ |
| 1:50 (low<br>mitochondrial<br>content) | ILMN1, ILMN6,<br>ILMN8, PACB2,<br>HYBR1,<br>HYBR2 | ILMN7 | ILMN2, ILMN4,<br>ILMN5,<br>PACB1, PACB3 | ILMN3 |
| 1:200 (high<br>mitochondrial<br>content) | PACB2,<br>HYBR1,<br>HYBR2 | ILMN6, ILMN7 | ILMN1, ILMN8,<br>PACB1, PACB3 | ILMN2, ILMN3,<br>ILMN4, ILMN5 |

<sup>\*</sup>The *de novo* assembly generated by each strategy was compared against the reference mitochondrial genome of *S. cerevisiae* using both QUAST and BLASTN. Unless a single contig was found to represent the complete mitochondrial genome, the assembled fraction of mitochondrial genome was determined based on the number of “missing reference bases” reported by QUAST, and further confirmed by the BLASTN result.

*Case Study III* (*Genomic architecture issue*). Lastly, we applied iWGS to three model eukaryotic genomes from different kingdoms and with different genomic architectures. Specifically, we analyzed *Drosophila melanogaster* (Adams, et al. 2000) and *Arabidopsis thaliana* (Arabidopsis Genome Initiative 2000), which are medium-sized animal and plant genomes, respectively, as well as *Plasmodium falciparum* 3D7 (Gardner, et al. 2002), a smaller protist genome with extremely low GC-content (∼19%). For all three genomes, the best assembly was generated by using only PacBio data (Table 3). In *D. melanogaster* and *A. thaliana*, several Illumina-only assemblies were of relatively high-quality (i.e. corrected scaffold N50 ≥ 100kb; Table S2), among which the best two were generated by ALLPATHS-LG and DISCOVAR (Table 3). However, all Illumina-only assemblies of *Pl. falciparum* 3D7 had considerably lower corrected scaffold N50 values, except for DISCOVAR whose sequencing strategy is unique in requiring a PE library with a limited insert size.

**Table 3.** Summary of top-ranking assemblies generated in case study III.

| Organism<br>(genome size) | Best assembly<br>from each<br>sequencing<br>strategy | Assembly statistics<sup>1</sup> |  |  |
| --- | --- | --- | --- | --- |
|  |  | Scaffold N50<br>(kb) | Largest scaffold<br>(kb) | Assembled<br>fraction of the<br>reference<br>genome |
| *D. melanogaster*<br>(137.55 Mb) | ILMN2 | 169.7 | 1007.9 | 89.1% |
|  | ILMN8 | 155.0 | 1007.7 | 91.8% |
|  | PACB2 | 5107.5 | 13108.3 | 99.3% |
|  | HYBR2 | 279.3 | 1536.8 | 89.7% |
| *A. thaliana*<br>(119.15 Mb) | ILMN2 | 307.0 | 1789.4 | 97.3% |
|  | ILMN8 | 266.6 | 2533.4 | 98.5% |
|  | PACB2 | 2065.7 | 8552.9 | 99.7% |
|  | HYBR2 | 289.3 | 1412.2 | 97.4% |
| *Pl. falciparum*<br>3D7<br>(23.29 Mb) | ILMN2 | 28.0 | 146.2 | 96.7% |
|  | ILMN8 | 222.0 | 729.9 | 98.4% |
|  | PACB2 | 282.9 | 1378.8 | 99.6% |
|  | HYBR2 | 15.5 | 91.5 | 97.4% |
| *Pl. falciparum*<br>IT<br>(real data) | ILMN2 | 167.1 | 641.7 | >100% |
|  | ILMN8 | 141.2 | 631.4 | 97.6% |
|  | PACB3 | 1574.7 | 3355.3 | 97.7% |
|  | HYBR2 | 198.4 | 602.1 | 92.2% |

<sup>1</sup>The statistics for simulation-based analysis of *D. melanogaster*, *A. thaliana*, and *Pl. falciparum* 3D7 are after correction for assembly errors using the reference genome, as reported by QUAST in GAGE mode. By default, QUAST (in GAGE mode) corrects contigs/scaffolds by breaking them at assembly errors larger than 5bp. The statistics for
real data based analysis of *Pl. falciparum* IT are calculated from the original *de novo* assemblies.

To examine how well the simulation-based predictions made by iWGS are supported by empirical data, we collected four real genome sequencing datasets from a previous study of *Pl. falciparum* IT [one overlapping 100bp PE library, one overlapping 250bp PE library, one MP library, and one PacBio library from (Otto, et al. 2014); Table S1] that were a good match to our simulated datasets, and ran the same set of assembly protocols. The best assembly was again generated by PacBio data alone, and the assemblies generated by ALLPATHS-LG, DISCOVAR (both are Illumina-only), and DBG2OLC (hybrid) were ranked next, while all other Illumina-only assembly protocols performed poorly (Table 3; Table S2). The results are largely consistent with our simulation study, suggesting that our simulation-based approach is indeed informative.

## Discussion

The design and analysis of *de novo* genome sequencing experiments is not trivial. On the design front, one has to balance between the complexity of the target genome, the strengths and weaknesses of each sequencing technology, and, importantly, the cost. Analysis is also challenging, as one is faced with multiple different algorithms and dozens of parameters. Although substantial efforts have been made to benchmark different approaches for genome assembly (Earl, et al. 2011; Salzberg, et al. 2012; Bradnam, et al. 2013; Magoc, et al. 2013), much less attention has been paid to investigating start-to-finish optimal sequencing strategies for a given genome [see (Chakraborty, et al. 2016) for one example].

iWGS is an automated tool that allows users to explicitly compare alternative experimental designs by using simulated sequencing data, even allowing users to estimate costs when these are known for the generation of each data type. We have illustrated the utility of iWGS in several case studies on mitochondrial and nuclear genomes with varying levels of complexity. For instance, our simulations suggest that Illumina-only sequencing strategies may be economical choices for the sequencing of relatively simple genomes (e.g. *Z. tritici*; Table S2), whereas PacBio data would be highly desirable for genomes of greater complexity (e.g. *Ps. fijiensis*, *C. geophilum*, and *Pl. falciparum*). Although not done here, iWGS could also be used to evaluate different combinations of sequencing assays (e.g. PE and MP libraries), read quality, read lengths, and sequencing depths. Empirical studies of both short- and long-read data have shown that these parameters are critical determinants of the quality of *de novo* genome assemblies (Utturkar, et al. 2014; Chakraborty, et al. 2016).

One key function of iWGS is the use of simulation data generated from a related reference genome to inform the experimental design for organisms lacking genomic data. A similar concept was previously used to evaluate sequencing strategies for cacao by using the rice genome as the reference (Haiminen, et al. 2011). In principle, one could apply iWGS on one or more related reference genomes that resemble the characteristics (e.g. genome size, repeat content, and sequence composition) of the sequencing target. However, if such reference genome is lacking, one solution is to start with a closely related reference genome and tune it toward the target (e.g. adjust GC- and repeat contents) by using third-party tools that simulate genome-wide evolution (Arenas and Posada 2014) before running iWGS. Alternatively, one may simply use iWGS with reference genomes that are of comparable complexity (e.g. similar in size and repeat content) regardless of the evolutionary relatedness. As suggested by previous studies, these factors not only influence the difficulty of genome assembly, but can also be excellent predictors of the assembly quality (Lee, et al. 2014). Therefore, iWGS could also be informative in evaluating the performance of alternative experimental designs on genomes with similar characteristics to the sequencing target.

Other important features of iWGS include the support for both Illumina short and PacBio long sequence reads and, correspondingly, a wide selection of software tools compatible with these data types, as well as the ability to analyze real data. In comparison, the support for third generation sequencing data is relatively limited in iMetAMOS and currently lacking in RAMPART. Given the increasing importance of long sequence reads in *de novo* genome assembly, iWGS aims to allow users to fully exploit the strength of long-read data and explore alternative ways of data analysis. Along these lines, several further developments can be envisioned. First, support for additional sequencing technologies, such as Oxford Nanopore, can be added as technologies become commercially available. In fact, the Celera Assembler, Canu, and SPAdes assemblers, which are supported by iWGS, can already utilize nanopore reads (Bankevich, et al. 2012; Berlin, et al. 2015). Similarly, realistic simulation of nanopore data will be possible once the patterns of errors and biases are better characterized using real data. Second, iWGS will continue to expand its functionality to achieve better assemblies. For instance, a number of assembly polishing tools can be integrated in iWGS to improve the quality of the final output, including Pilon (Walker, et al. 2014), Quiver (Chin, et al. 2013), and Nanopolish (Loman, et al. 2015) which use Illumina, PacBio, and nanopore data, respectively. In addition, iWGS currently uses Metassembler for meta-assembly; in the future, other meta-assembly tools that support assemblies based on PacBio data alone, such as quickmerge (Chakraborty, et al. 2016), could be added. Lastly, it would be beneficial to enable users to add new software tools to iWGS in order to stay up-to-date with the rapid advances in genome assembly and other aspects of HTS data analysis. We intend to provide periodic updates, and the expert user can edit iWGS on their own. In summary, iWGS is a flexible, expandable, and easy to use pipeline that will aid in the design and execution of genome assembly experiments across the tree of life.

## Acknowledgements

We thank Francis Martin for providing access to unpublished genome data for the *Cenococcum geophilum* genome produced by the U.S. Department of Energy Joint Genome Institute, a DOE Office of Science User Facility, supported by the Office of Science of the U.S. Department of Energy under Contract No. DE-AC02-05CH11231. We thank Branden Timm for technical support on the WEI cluster. This work was conducted in part using the resources of the Advanced Computing Center for Research and Education at Vanderbilt University and the Wisconsin Energy Institute (WEI) cluster at the University of Wisconsin-Madison. This work was funded by the National Science Foundation (DEB-1442113 to AR; DEB-1253634 and DEB-1442148 to CTH), in part by the DOE Great Lakes Bioenergy Research Center (DOE Office of Science BER DEFC02- 07ER64494 to CTH), the USDA National Institute of Food and Agriculture (Hatch Project 1003258 to CTH), and the National Institutes of Health (NIAID AI105619 to AR). CTH is an Alfred Toepfer Faculty Fellow, supported by the Alexander von Humboldt Foundation. CTH is a Pew Scholar in the Biomedical Sciences, supported by the Pew Charitable Trusts. Mention of trade names or commercial products in this publication is solely for the purpose of providing specific information and does not imply recommendation or endorsement by the U.S. Department of Agriculture. USDA is an equal opportunity provider and employer.

## References

Abbas MM, Malluhi QM, Balakrishnan P. 2014. Assessment of de novo assemblers for draft genomes: a case study with fungal genomes. BMC Genomics. 15 Suppl 9:S10.

Adams MD, Celniker SE, Holt RA, Evans CA, Gocayne JD, Amanatides PG, Scherer SE, Li PW, Hoskins RA, Galle RF, et al. 2000. The genome sequence of Drosophila melanogaster. Science. 287:2185–2195.

Arabidopsis Genome Initiative. 2000. Analysis of the genome sequence of the flowering plant Arabidopsis thaliana. Nature. 408:796–815.

Arenas M, Posada D. 2014. Simulation of genome-wide evolution under heterogeneous substitution models and complex multispecies coalescent histories. Mol Biol Evol. 31:1295–1301.

Baker E, Wang B, Bellora N, Peris D, Hulfachor AB, Koshalek JA, Adams M, Libkind D, Hittinger CT. 2015. The Genome Sequence of Saccharomyces eubayanus and the Domestication of Lager-Brewing Yeasts. Mol Biol Evol. 32:2818–2831.

Bankevich A, Nurk S, Antipov D, Gurevich AA, Dvorkin M, Kulikov AS, Lesin VM, Nikolenko SI, Pham S, Prjibelski AD, et al. 2012. SPAdes: a new genome assembly algorithm and its applications to single-cell sequencing. J Comput Biol. 19:455–477.

Berlin K, Koren S, Chin CS, Drake JP, Landolin JM, Phillippy AM. 2015. Assembling large genomes with single-molecule sequencing and locality-sensitive hashing. Nat Biotechnol. 33:623–630.

Bradnam KR, Fass JN, Alexandrov A, Baranay P, Bechner M, Birol I, Boisvert S, Chapman JA, Chapuis G, Chikhi R, et al. 2013. Assemblathon 2: evaluating de novo methods of genome assembly in three vertebrate species. Gigascience. 2:10.

Chakraborty M, Baldwin-Brown JG, Long AD, Emerson JJ. 2016. Contiguous and accurate de novo assembly of metazoan genomes with modest long read coverage. bioRxiv.

Chapman JA, Ho I, Sunkara S, Luo S, Schroth GP, Rokhsar DS. 2011. Meraculous: de novo genome assembly with short paired-end reads. PLoS One. 6:e23501.

Chikhi R, Medvedev P. 2014. Informed and automated k-mer size selection for genome assembly. Bioinformatics. 30:31–37.

Chin CS, Alexander DH, Marks P, Klammer AA, Drake J, Heiner C, Clum A, Copeland A, Huddleston J, Eichler EE, et al. 2013. Nonhybrid, finished microbial genome assemblies from long-read SMRT sequencing data. Nat Methods. 10:563–569.

Denton JF, Lugo-Martinez J, Tucker AE, Schrider DR, Warren WC, Hahn MW. 2014. Extensive error in the number of genes inferred from draft genome assemblies. PLoS Comput Biol. 10:e1003998.

Earl D, Bradnam K, St John J, Darling A, Lin D, Fass J, Yu HO, Buffalo V, Zerbino DR, Diekhans M, et al. 2011. Assemblathon 1: a competitive assessment of de novo short read assembly methods. Genome Res. 21:2224–2241.

Foury F, Roganti T, Lecrenier N, Purnelle B. 1998. The complete sequence of the mitochondrial genome of Saccharomyces cerevisiae. FEBS Lett. 440:325–331.

Freel KC, Friedrich A, Schacherer J. 2015. Mitochondrial genome evolution in yeasts: an all-encompassing view. FEMS Yeast Res. 15:fov023.

Gardner MJ, Hall N, Fung E, White O, Berriman M, Hyman RW, Carlton JM, Pain A, Nelson KE, Bowman S, et al. 2002. Genome sequence of the human malaria parasite Plasmodium falciparum. Nature. 419:498–511.

Genome 10K Community of Scientists. 2009. Genome 10K: a proposal to obtain whole-genome sequence for 10,000 vertebrate species. J Hered. 100:659–674.

Glenn TC. 2011. Field guide to next-generation DNA sequencers. Mol Ecol Resour. 11:759–769.

Gnerre S, Maccallum I, Przybylski D, Ribeiro FJ, Burton JN, Walker BJ, Sharpe T, Hall G, Shea TP, Sykes S, et al. 2011. High-quality draft assemblies of mammalian genomes from massively parallel sequence data. Proc Natl Acad Sci U S A. 108:1513–1518.

Goodwin SB, M’Barek S B, Dhillon B, Wittenberg AH, Crane CF, Hane JK, Foster AJ, Van der Lee TA, Grimwood J, Aerts A, et al. 2011. Finished genome of the fungal wheat pathogen Mycosphaerella graminicola reveals dispensome structure, chromosome plasticity, and stealth pathogenesis. PLoS Genet. 7:e1002070.

Grigoriev IV, Cullen D, Goodwin SB, Hibbett D, Jeffries TW, Kubicek CP, Kuske C, Magnuson JK, Martin F, Spatafora JW, et al. 2011. Fueling the future with fungal genomics. Mycology. 2:192–209.

Gurevich A, Saveliev V, Vyahhi N, Tesler G. 2013. QUAST: quality assessment tool for genome assemblies. Bioinformatics. 29:1072–1075.

Haiminen N, Kuhn DN, Parida L, Rigoutsos I. 2011. Evaluation of methods for de novo genome assembly from high-throughput sequencing reads reveals dependencies that affect the quality of the results. PLoS One. 6:e24182.

Hittinger CT, Rokas A, Bai FY, Boekhout T, Goncalves P, Jeffries TW, Kominek J, Lachance MA, Libkind D, Rosa CA, et al. 2015. Genomics and the making of yeast biodiversity. Curr Opin Genet Dev. 35:100–109.

Hu X, Yuan J, Shi Y, Lu J, Liu B, Li Z, Chen Y, Mu D, Zhang H, Li N, et al. 2012. pIRS: Profile-based Illumina pair-end reads simulator. Bioinformatics. 28:1533–1535.

Huang W, Li L, Myers JR, Marth GT. 2012. ART: a next-generation sequencing read simulator. Bioinformatics. 28:593–594.

Hunt M, Kikuchi T, Sanders M, Newbold C, Berriman M, Otto TD. 2013. REAPR: a universal tool for genome assembly evaluation. Genome Biol. 14:R47.

Kajitani R, Toshimoto K, Noguchi H, Toyoda A, Ogura Y, Okuno M, Yabana M, Harada M, Nagayasu E, Maruyama H, et al. 2014. Efficient de novo assembly of highly heterozygous genomes from whole-genome shotgun short reads. Genome Res. 24:1384–1395.

Koren S, Harhay GP, Smith TP, Bono JL, Harhay DM, McVey SD, Radune D, Bergman NH, Phillippy AM. 2013. Reducing assembly complexity of microbial genomes with single-molecule sequencing. Genome Biol. 14:R101.

Koren S, Phillippy AM. 2015. One chromosome, one contig: complete microbial genomes from long-read sequencing and assembly. Curr Opin Microbiol. 23:110–120.

Koren S, Treangen TJ, Hill CM, Pop M, Phillippy AM. 2014. Automated ensemble assembly and validation of microbial genomes. BMC Bioinformatics. 15:126.

Kumar S, Subramanian S. 2002. Mutation rates in mammalian genomes. Proc Natl Acad Sci U S A. 99:803–808.

Lee H, Gurtowski J, Yoo S, Marcus S, McCombie WR, Schatz M. 2014. Error correction and assembly complexity of single molecule sequencing reads. bioRxiv.

Loman NJ, Quick J, Simpson JT. 2015. A complete bacterial genome assembled de novo using only nanopore sequencing data. Nat Methods. 12:733–735.

Luo R, Liu B, Xie Y, Li Z, Huang W, Yuan J, He G, Chen Y, Pan Q, Liu Y, et al. 2012. SOAPdenovo2: an empirically improved memory-efficient short-read de novo assembler. Gigascience. 1:18.

Magoc T, Pabinger S, Canzar S, Liu X, Su Q, Puiu D, Tallon LJ, Salzberg SL. 2013. GAGE-B: an evaluation of genome assemblers for bacterial organisms. Bioinformatics. 29:1718–1725.

Mapleson D, Drou N, Swarbreck D. 2015. RAMPART: a workflow management system for de novo genome assembly. Bioinformatics. 31:1824–1826.

McIlwain SJ, Peris D, Sardi M, Moskvin OV, Zhan F, Myers KS, Riley NM, Buzzell A, Parreiras LS, Ong IM, et al. 2016. Genome Sequence and Analysis of a Stress-Tolerant, Wild-Derived Strain of Saccharomyces cerevisiae Used in Biofuels Research. G3 (Bethesda). 6:1757–1766.

Mewes HW, Albermann K, Bahr M, Frishman D, Gleissner A, Hani J, Heumann K, Kleine K, Maierl A, Oliver SG, et al. 1997. Overview of the yeast genome. Nature. 387:7–65.

Myers EW, Sutton GG, Delcher AL, Dew IM, Fasulo DP, Flanigan MJ, Kravitz SA, Mobarry CM, Reinert KH, Remington KA, et al. 2000. A whole-genome assembly of Drosophila. Science. 287:2196–2204.

Nagarajan N, Pop M. 2013. Sequence assembly demystified. Nat Rev Genet. 14:157–167.

Ohm RA, Feau N, Henrissat B, Schoch CL, Horwitz BA, Barry KW, Condon BJ, Copeland AC, Dhillon B, Glaser F, et al. 2012. Diverse lifestyles and strategies of plant pathogenesis encoded in the genomes of eighteen Dothideomycetes fungi. PLoS Pathog. 8:e1003037.

Ono Y, Asai K, Hamada M. 2013. PBSIM: PacBio reads simulator--toward accurate genome assembly. Bioinformatics. 29:119–121.

Otto TD, Rayner JC, Bohme U, Pain A, Spottiswoode N, Sanders M, Quail M, Ollomo B, Renaud F, Thomas AW, et al. 2014. Genome sequencing of chimpanzee malaria parasites reveals possible pathways of adaptation to human hosts. Nat Commun. 5:4754.

Reddy TB, Thomas AD, Stamatis D, Bertsch J, Isbandi M, Jansson J, Mallajosyula J, Pagani I, Lobos EA, Kyrpides NC. 2015. The Genomes OnLine Database (GOLD) v.5: a metadata management system based on a four level (meta)genome project classification. Nucleic Acids Res. 43:D1099–1106.

Ribeiro FJ, Przybylski D, Yin S, Sharpe T, Gnerre S, Abouelleil A, Berlin AM, Montmayeur A, Shea TP, Walker BJ, et al. 2012. Finished bacterial genomes from shotgun sequence data. Genome Res. 22:2270–2277.

Robinson GE, Hackett KJ, Purcell-Miramontes M., Brown SJ, Evans JD, Goldsmith MR, Lawson D, Okamuro J, Robertson HM, Schneider DJ. 2011. Creating a buzz about insect genomes. Science. 331:1386.

Rokas A, Abbot P. 2009. Harnessing genomics for evolutionary insights. Trends Ecol Evol. 24:192–200.

Rokas A, Williams BL, King N, Carroll SB. 2003. Genome-scale approaches to resolving incongruence in molecular phylogenies. Nature. 425:798–804.

Ross MG, Russ C, Costello M, Hollinger A, Lennon NJ, Hegarty R, Nusbaum C, Jaffe DB. 2013. Characterizing and measuring bias in sequence data. Genome Biol. 14:R51.

Roux J, Privman E, Moretti S, Daub JT, Robinson-Rechavi M., Keller L. 2014. Patterns of positive selection in seven ant genomes. Mol Biol Evol. 31:1661–1685.

Salichos L, Rokas A. 2013. Inferring ancient divergences requires genes with strong phylogenetic signals. Nature. 497:327–331.

Salikhov K, Sacomoto G, Kucherov G. 2013. Using Cascading Bloom Filters to Improve the Memory Usage for de Brujin Graphs. In: Darling A, Stoye J, editors. Algorithms in Bioinformatics: Springer Berlin Heidelberg. p. 364–376.

Salzberg SL, Phillippy AM, Zimin A, Puiu D, Magoc T, Koren S, Treangen TJ, Schatz MC, Delcher AL, Roberts M, et al. 2012. GAGE: A critical evaluation of genome assemblies and assembly algorithms. Genome Res. 22:557–567.

Simpson JT, Durbin R. 2012. Efficient de novo assembly of large genomes using compressed data structures. Genome Res. 22:549–556.

Simpson JT, Pop M. 2015. The Theory and Practice of Genome Sequence Assembly. Annu Rev Genomics Hum Genet. 16:153–172.

Simpson JT, Wong K, Jackman SD, Schein JE, Jones SJ, Birol I. 2009. ABySS: a parallel assembler for short read sequence data. Genome Res. 19:1117–1123.

Solieri L. 2010. Mitochondrial inheritance in budding yeasts: towards an integrated understanding. Trends Microbiol. 18:521–530.

Utturkar SM, Klingeman DM, Land ML, Schadt CW, Doktycz MJ, Pelletier DA, Brown SD. 2014. Evaluation and validation of de novo and hybrid assembly techniques to derive high-quality genome sequences. Bioinformatics. 30:2709–2716.

Walker BJ, Abeel T, Shea T, Priest M, Abouelliel A, Sakthikumar S, Cuomo CA, Zeng Q, Wortman J, Young SK, et al. 2014. Pilon: an integrated tool for comprehensive microbial variant detection and genome assembly improvement. PLoS One. 9:e112963.

Weisenfeld NI, Yin S, Sharpe T, Lau B, Hegarty R, Holmes L, Sogoloff B, Tabbaa D, Williams L, Russ C, et al. 2014. Comprehensive variation discovery in single human genomes. Nat Genet. 46:1350–1355.

Wences AH, Schatz MC. 2015. Metassembler: merging and optimizing de novo genome assemblies. Genome Biol. 16:207.

Ye C, Hill C, Wu S, Ruan J, Zhanshan, Ma. 2014. DBG2OLC: Efficient Assembly of Large Genomes Using Long Erroneous Reads of the Third Generation Sequencing Technologies. ArXiv e-prints.

Ye C, Ma ZS, Cannon CH, Pop M, Yu DW. 2012. Exploiting sparseness in de novo genome assembly. BMC Bioinformatics. 13 Suppl 6:S1.

Zerbino DR, Birney E. 2008. Velvet: algorithms for de novo short read assembly using de Bruijn graphs. Genome Res. 18:821–829.

Zhang G, Li C, Li Q, Li B, Larkin DM, Lee C, Storz JF, Antunes A, Greenwold MJ, Meredith RW, et al. 2014. Comparative genomics reveals insights into avian genome evolution and adaptation. Science. 346:1311–1320.

Zhou X, Rokas A. 2014. Prevention, diagnosis and treatment of high-throughput sequencing data pathologies. Mol Ecol. 23:1679–1700.

Zimin AV, Marcais G, Puiu D, Roberts M, Salzberg SL, Yorke JA. 2013. The MaSuRCA genome assembler. Bioinformatics. 29:2669–2677.

